# FGF2 modulates simultaneously the mode, the rate of division and the growth fraction in cultures of Radial Glia

**DOI:** 10.1101/707463

**Authors:** Mario Ledesma-Terrón, Nuria Peralta-Cañadas, David G. Míguez

## Abstract

Radial Glial progenitors in the mammalian developing neocortex have been shown to follow a deterministic differentiation program restricted to an asymmetric-only mode of division. This feature seems incompatible with their well known ability to expand in number when cultured *in vitro*, driven by Fibroblast Growth Factor 2 and other mitogenic signals. The changes in their differentiation dynamics that allow this transition from *in vivo* asymmetric-only division mode to an *in vitro* self-renewing culture have not been fully characterized. Here we combine experiments of Radial Glia cultures with theory and numerical models to show that Fibroblast Growth Factor 2 has a triple effect by simultaneously increasing the growth fraction, promoting symmetric divisions and shortening the length of the cell cycle. This combined effect of Fibroblast Growth Factor 2 in the differentiation dynamics of Radial Glial progenitors partner to establish and sustain a pool of rapidly proliferating *in vitro* pool of Radial Glial progenitors.

## INTRODUCTION

The neocortex constitutes the main part of the mammalian brain, and the location where the processing of all higher-order brain functions resides. Understanding its formation is one of the major interests in the field of Developmental Biology (Lodato and Arlotta, 2015). The neocortex develops from a stratified neuroepithelium, called the neural tube, into a complex structure of six horizontal layers of excitatory and inhibitory neurons (Matsuzaki and Shitamukai, 2015). Neurogenesis in the developing neocortex initiates when self-renewing neuroepithelial progenitors (NEP) transform into apical and basal Radial Glial (RG) progenitor cells and start to produce neurons and intermediate neuronal precursors (Beattie and Hippenmeyer, 2017; Taverna et al., 2014). Since the discovery that RG constitute the progenitors of potentially all neurons in the vertebrate neocortex (Frederiksen and McKay, 1988; Hartfuss et al., 2001; Miyata et al., 2001; Noctor et al., 2004), a great effort has been focused in identifying their features and properties: how they coordinate in time and space to form the multiple layers of the neocortex?; which signals control their fate?; and how these signals orchestrate the correct balance between proliferation or differentiation during neurogenesis?.

In principle, this balance can be robustly achieved via stochastic or deterministic cell decisions (Losick and Desplan, 2008). In brief, stochastic models assume certain probability of differentiation that depends on the intracellular and extracellular signals that the cell is receiving. In this context, the fate at the single cell level is unpredictable and the balance between proliferation and differentiation is regulated at the level of the population (Teles et al., 2013). On the other hand, deterministic models of stem cell differentiation assume that the fate of the progeny is fixed and, therefore, the correct balance between the numbers of different types of neurons is achieved at the single cell level (Müller-Sieburg et al., 2002).

The dynamics of differentiation is often characterized based on the fate of the two daughter cells of a cell division relative to each other (Kosodo et al., 2004). This way, proliferating progenitors can perform *pp* (progenitor-progenitor), *pd* (progenitor-differentiated) and *dd* (differentiated-differentiated) divisions (Huttner and Kosodo, 2005).

In this context, differentiation in the developing chick spinal cord (Míguez, 2015), in the zebrafish retina (He et al., 2012; Chen et al., 2012), epidermis (Clayton et al., 2007), airway epithelium (Teixeira et al., 2013), germline (Klein et al., 2010), and the intestine (Snippert et al., 2010) of mice follow an stochastic model. In these systems, progenitors can potentially perform each of the three types of division, and the corresponding rates are probabilistic and change overtime. On the other hand, the differentiation of RG in the mammalian brain has been shown to follow a deterministic asymmetric-only mode of division (Gao et al., 2014; Beattie and Hippenmeyer, 2017).

Several years ago, the group of Austin Smith showed that RG extracted from mouse developing neocortex can be succesfully cultured *in vitro* (Conti et al., 2005). Driven by the multiple phenotypic similarities between neuronal precursors differentiated from embryonic stems cells in culture and RG, authors suggested that these neuronal precursors are the culture analogs to RG. In the same paper and driven by this observation, they also showed that *in vitro* cultures of RG can be established with Fibroblast Growth Factor 2 (FGF2) as the key mitogen that facilitates their expansion (Conti et al., 2005).

FGF2 is an extensively studied neurogenic factor for proliferation and differentiation of multipotent neural stem cells both during development and in the adult mouse brain (Kang and Hébert, 2015). FGF2 has been shown to be necessary for cell proliferation and neurogenesis *in vivo*, and to induce additional mitoses in progenitor cells *in vitro* (Raballo et al., 2000). In addition, stem cells from the adult mouse brain have been shown to proliferate and self-renew *in vitro* in the presence of FGF2 (Gritti et al., 1996). On the other hand, FGF2 stimulation have been shown to control the fate, migration and differentiation but not the proliferation of neuronal progenitors *in vivo* (Dono et al., 1998), while more recent studies do show an impact in promoting the cell cycle progression in cultures of Rat glioblastoma cells (Baguma-Nibasheka et al., 2012).

From all these potential effects of FGF2, the specific features that facilitate the transition of RG from a non-expanding population *in vivo* that can only perform asymmetric *pd* divisions (and therefore, incompatible with cell expansion in number), to a self-renewing *in vitro* culture have not been quantitatively characterized in detail. In principle, this transition can be achieved by reducing the rate of neurogenesis, by promoting proliferative (at the expenses of asymmetric or symmetric differentiative) divisions, by increasing the proliferation rate (by shortening the cell cycle), by inducing cell cycle reentry of quiescent progenitors (i.e., increasing the growth fraction), by reducing apoptosis (as a pro-survival signal), inducing intermediate progenitors (that perform additional terminal divisions), or also by shifting RG towards its less mature NEP phenotype (that perform pp divisions *in vivo*).

In this paper, we quantify the specific effects of FGF2 on key features of the proliferation and differentiation dynamics of RG that allow them to be cultured and expanded *in vitro*. To to that, we quantify values of cell numbers of RG and differentiated neurons extracted from mouse developing cerebral cortex and cultured in the presence of different FGF2 concentrations and at different time points. These values inform a theoretical framework based on a branching process formalism (Míguez, 2015) that provides average values of mode and rate of division of the RG population with temporal resolution. Our results show that FGF2 does not affect the rate of neurogenesis (i.e., the amount of differentiated neurons produced), it does not promote NEP or intermediate progenitor phenotype and it does not affect significantly the apoptosis rate. On the other hand, FGF2 does promote symmetric *pp* divisions, it increases the growth fraction, and shortens the average cell cycle length. These three key effects when combined, strongly facilitate the propagation and expansion of the culture.

In addition, discrepancies between predictions for the cell cycle length and growth fraction using several methods in our study pointed us to compare the accuracy of several common methodologies used to measure cell cycle features. To do that, we use a numerical model to show that methods based on cumulative thymidine analogs (such as Edu and BrdU) are not accurate in conditions of variable differentiation dynamics. On the other hand, the method based on branching process formalism performs better when mode and/or rate of division are changing, which is the case in our RG cultures and many other *in vivo* developmental systems. In addition, the branching process method is superior due to its temporal resolution, robustness, minimal interference with cell homeostasis, and simplicity of use.

## Results

### FGF2 stimulation increases the growth rate of cultures of RG by shortening the length of the the cell cycle

To initially test how the dynamics of growth and differentiation of RG *in vitro* is modulated by FGF2, cells derived from the developing neocortex of mouse embryos at E11-11.5 are extracted, plated and cultured following standard protocols (Hilgenberg and Smith, 2007). Starting at 24 hours post plating (hpp), samples are then fixed at three different time points and stained with Hoechst (Fig. 1A). Quantification of the number of cells in a field of view of fixed dimensions using an automated segmentation tool developed in house (see Supplementary Methods) is shown in Fig. 1B for two culture conditions: SC and SC+FGF, where the standard culture media is supplemented with an increased concentration of FGF2 ligand (see Methods). In both conditions, the number of cells increases, but the growth is only statistically significant (*P* < 0.05) in SC+FGF conditions.

**Fig. 1.**
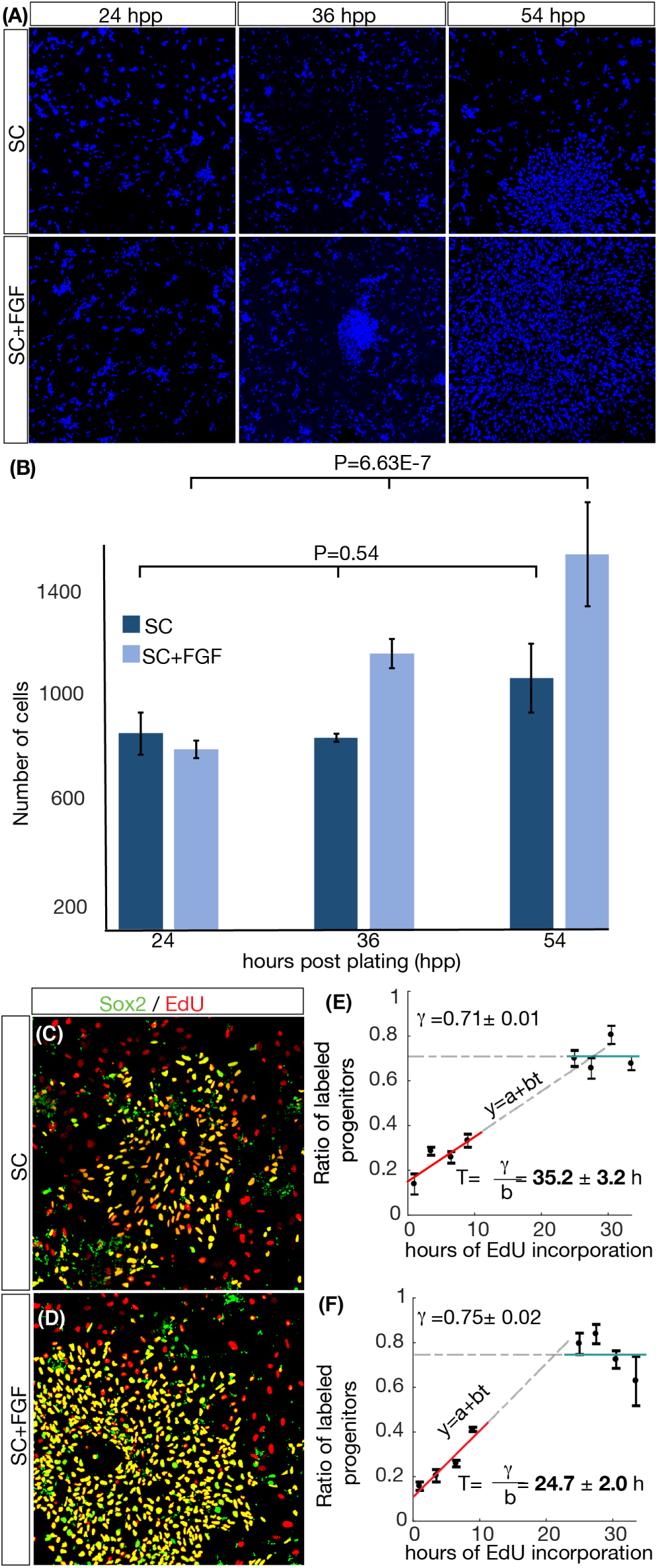
FGF2 shortens the division time of cycling RG *in vitro*. (A) Snapshots of RG cultures at different hours post plating (hpp) stained with Hoechst and growing at SC and SC+FGF culture conditions. (B) Total cell numbers in a field of view of 0.6 mm × 0.6 mm at different time points. Error bars correspond to standard error of the mean value between multiple samples of similar conditions. (C-D) Sox2 (green) and and EdU (red) staining to mark progenitors that have gone through S-Phase in 24 hours of EdU incorporation. (E-F) Cumulative curve of EdU positive progenitors shows that cells in SC+FGF conditions cycle faster (T= 24.7 ± 2.0 hours) than in SC (T=35.2 ± 3.5 hours), while the growth fraction *γ* remains similar.

To study in detail how FGF2 affects the length of the cell cycle of the cycling progenitors, we perform 5-ethynyl-2’-deoxyuridine (EdU) cumulative labeling experiments to measure changes in the length of the average cell cycle. BrdU (Nowakowski et al., 1989), EdU (Salic and Mitchison, 2008; Buck et al., 2008) and other thymidine analogs constitute the most used tool to estimate the cell cycle length of cells in many contexts (Alexiades and Cepko, 1996). The methodology is based on the replacement of endogenous thymidine during DNA synthesis with traceable compounds (Takahashi, 1966; Takahashi et al., 1996). The length of the average cell cycle is then inferred from the dynamics of the incorporation of these compounds into the DNA of cycling cells (Macdonald, 1970).

To estimate the average cell cycle length of the population, samples are cultured in the presence of EdU and then fixed at different time points (corresponding to different times of EdU incorporation). Combined nuclear Hoechst staining with EdU detection assay and immunostaining against Sox2 is used to identify all progenitors that have passed through S-phase for each EdU incubation time.

The cell cycle length *T* and the growth fraction *γ* are calculated using the standard cumulative curve methodology based on linear regression (see Methods). Representative snapshots are shown in Fig. 1C-F. The resulting cumulative curves (Figs. 1E, F) reveal that *γ* remains at around 72% for both conditions tested, while *T* depends strongly on the culture conditions (T=35.2 ± 3.5 hours for SC, T= 24.7 ± 2.0 hours for SC+FGF). In conclusion, our results show that FGF2 stimulation shortens the average cell cycle length in cultures of RG *in vitro*, while its effect in the growth fraction is not statistically significant.

### FGF2 stimulates the generation of progenitors in culture

The previous section shows that FGF2 affects the rate of division. To study the effect of FGF2 in the number of cells of each specific population of RG progenitors and differentiated neurons, we extract the neocortex of mouse embryos at E11-11.5 and plate cells at same initial cell density in different wells. Next, cells are cultured under the two conditions of FGF2 and samples are then fixed them every 2-4 hours, starting at 24 hours post plating (hpp). Next, samples are stained using antibodies against Sox2 and Map2 to identify progenitors and differentiated cells, respectively. We then identify the fate of each cell based on the intensity of Sox2 and Map2 staining using our segmentation framework in images of 0.6 mm × 0.6 mm (see Supplementary Methods).

Results are shown in Fig. 2 for the two conditions tested: SC and SC+FGF. Output provided by the segmentation script is plotted in Figs 2B, C. Assuming the typical logistic growth model (Juarez et al., 2016) for proliferating cells in cultures, the corresponding sigmoidal curve fitting is also plotted (green, red, and blue lines for RG, neurons and total cells, respectively). The data shows that an initial regime of reduced change in cell numbers is followed by an increase in both cell types until the system reaches a regime where few new cells are being generated. In both conditions, the amount of progenitors (green data points, green line) and differentiated cells (red data points, red line) increases with statistical significance (*P* < 0.05) but the increase in progenitors is statistically more significant in conditions of SC+FGF (P= 7.25E-09) that in SC conditions (P= 7.60E-03).

**Fig. 2.**
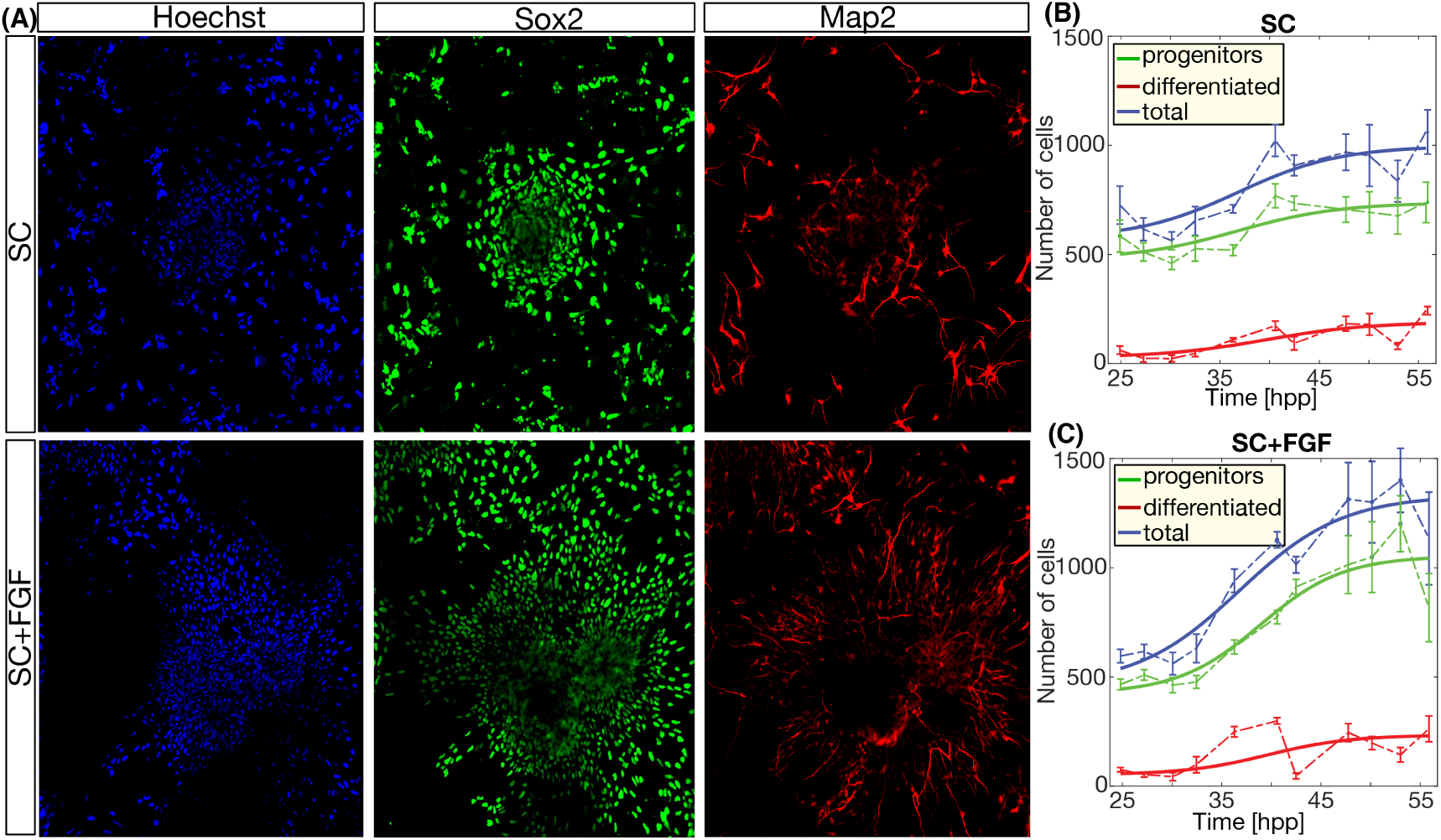
FGF stimulation increases the amount of progenitor cells. (A) Snapshots of RG cultures at 24 hours post plating showing nuclei (Hoechst), progenitors (stained with Sox2) and differentiated neurons (stained with Map2). (B-C) Quantification of the number of cells of each type in both culture conditions at different time points shows an increased number of progenitors is SC+FGF, compared to SC conditions. Error bars correspond to standard error of the mean. Lines correspond to nonlinear sigmoidal fitting of the experimental data points.

In principle, this increase in the progenitor population could be explained by an increase in the population of intermediate progenitors (Molyneaux et al., 2007). This sub-population of cycling cells emerge from asymmetric division of the RG *in vivo*, and they are able to perform a terminal division to produce two terminally differentiated neurons (Hutton and Pevny, 2011). Immunofluorescence against Tbr2, a marker for intermediate progenitors shows no Tbr2 positive cells in the two culture conditions tested (data not shown). This is in agreement with the effect of FGF2 in inhibiting the transition from RG to intermediate progenitor (Kang et al., 2009) (FGF2 is in the culture media in both experimental conditions: SC and SC+FGF).

Another possibility that could explain this increase in the number of RG progenitors is the presence of neuroepithelial progenitors (NEPs) in the culture (that have been shown to proliferate *in vivo* via *pp* divisions) (Beattie and Hippenmeyer, 2017; Taverna et al., 2014). Quantification of immunofluorescence against Pax6, a well characterized marker for RG (Suter et al., 2009) that is not present in NEPs (Elsen et al., 2018) shows that close to 100% of all Sox2 positive progenitors are also positive for Pax6 (Supplementary Figure S1A), suggesting that FGF2 stimulation does not result in the presence of neuroepithelial progenitors.

In conclusion, the increase in FGF2 concentration does not produce intermediate or NEP progenitors, and results in more RG and similar number of differentiated cells, showing that the population of cycling progenitors does not grow at the expense of the terminally differentiated cells.

### Branching process formalism predicts variable mode of division that is affected by FGF2 stimulation

The previous observation suggests that, apart from the changes in the cell cycle length, FGF2 may also be affecting the mode of division of the RG. It has been shown previously that the fate of differentiating RG can be modulated by FGF2, by changing the differentiation progeny of RG from neurons to glia (Qian et al., 1997). To quantify the effect of FGF2 in the mode of division, we take advantage of a branching process theoretical formalism developed by our lab (Míguez, 2015). In brief, the tool provides the average rates of each mode of division with temporal resolution simply based on numbers of progenitors and differentiated cells at different time points (see Supplementary Methods).

Input data of the framework are the numbers of progenitors and differentiated cells, the rate of apoptosis and the growth fraction. To obtain the average rate of apoptosis, we perform immunostaining against anti-Cleaved Caspase 3 at three time points in the cultures at SC and SC+FGF conditions. Comparison between both conditions show a very reduced rate of apoptosis that is not significantly affected by the addition of extra FGF2 (Supplementary Figure S1B).

Next, the fitted values of cell numbers for progenitors and differentiated cells from the previous section, and the apoptosis rate are used as input in Eq. 1 in the Supplementary Methods (Míguez, 2015) to obtain the average value of *pp-dd* divisions. Results are shown in Fig. 3A. In both cases, differentiation appears to increase in time, and this change is reduced in SC-FGF conditions. Interestingly, both situations show values of *pp* − *dd* ≠ 0, which would correspond to the *in vivo* situation of asymmetric only divisions *pd* = 1 (*pp* + *pd* + *dd* = 1). Also, the average rate of differentiation is not constant in time, with the maximum change in the differentiation dynamics occurs around 36-37 hpp. Comparison between the two curves show that the value of *pp* − *dd* predicted is higher when more FGF2 is present in the culture media, which corresponds with the higher increase in the number of progenitors observed in SC-FGF conditions (Fig. 2C).

**Fig. 3.**
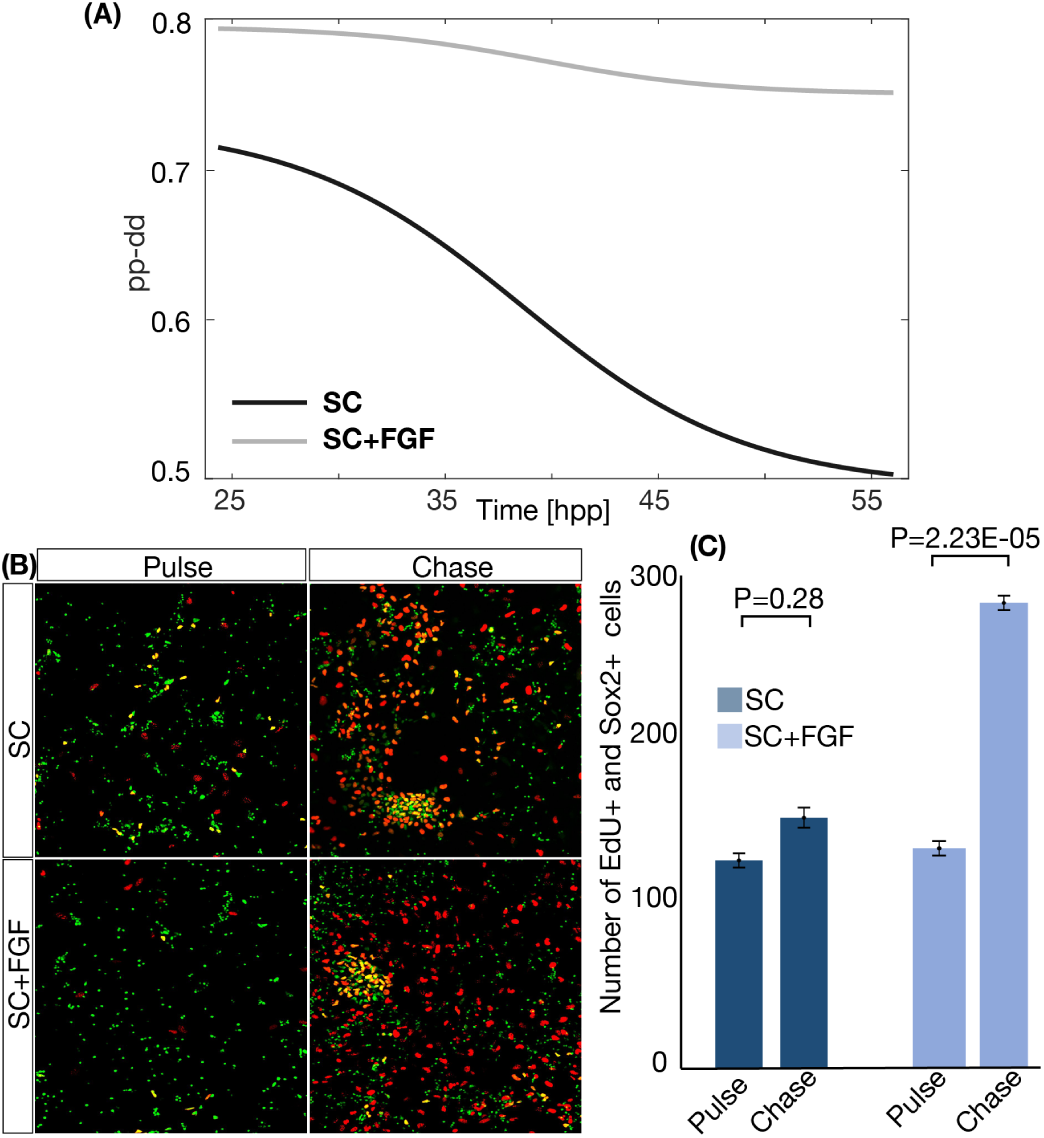
FGF affects the proportion of symmetric proliferative divisions in RG culture. (A) Plot of the average value of *pp* − *dd* of the population of RG under SC (black) and SC+FGF (grey) conditions. (B) Representative images showing Sox2 and EdU (stained in green and red respectively) for “pulse” and “chase” time points. (C) Quantification of the number of Sox2 and EdU positive cells for time-points for SC and SC+FGF conditions. Error bars correspond to standard error of the mean value between independent repeats of the experiment.

To further validate the result that an increase in FGF2 increases the amount of pp divisions, we designed an experiment based on Pulse-and-Chase of EdU labelled cells. Do do that, we plate cells from mouse developing neocortex following the procedure explained in Methods section. Next, cells are cultured in SC and SC+FGF conditions until 33 hpp. At this point, a pulse of 30-minutes of EdU is applied to all samples. A number of samples are fixed at this time point (and labeled as “Pulse” time point). The rest of samples are washed with fresh culture media 5 times to remove the Edu (see Methods). These samples are cultured for another 15 hours (corresponding to the predicted average *T* for SC+FGF conditions during this time, to ensure that labeled cells cannot cycle more than once in any of the culture conditions). Next, cells are fixed and stained with Hoechst, EdU and Sox2 immunostaining. Finally, the number of Sox2+/EdU+ cells at the time of the pulse (33 hpp) and chase (47 hpp) is quantified using our automated image analysis tool (see Supplementary Methods). Results are shown in Fig. 3B-C. The number of progenitors labeled with EdU does not change significantly in SC conditions, consistent with a large proportion of asymmetric divisions (i.e, one EdU+ RG produces two EdU+ cells: one RG and one neuron, so the amount of EdU+ RG remains constant) or a balanced ratio between *pp* and *dd*. On the other hand, in conditions of SC+FGF, we see a statistically significant (P<0.05) increase in the number of EdU+ RG when comparing “pulse” and “chase” time points. This result shows that more RG originally labeled with the short EdU pulse, divided and produced more RG when FGF2 is increased.

### The length of the cell cycle is variable and shortens in conditions of FGF2 stimulation

The Branching Process formalism also provides the average cell cycle length of the progenitors in the culture with temporal resolution (Eq. 2 in Supplementary Material). This equation uses as additional input the value of the growth fraction *γ*, which can be indirectly obtained from the EdU experiments in Figs. 1E-F. To obtain a more direct estimation of the amount of quiescent progenitors, we perform immunofluorescence against KI67 at different time points (Fig. 4A) (Scholzen and Gerdes, 2000). The automated quantification of the number of Sox2+ cells that are also KI67+ in both culture conditions (Fig. 4B) shows statistically significant differences between SC and SC+FGF conditions, contrary to the results obtained with EdU cumulative curves in Figs. 1E-F. In SC conditions, the growth fraction is around 55%, while the value in SC+FGF conditions is closer to 90%. This discrepancy between the EdU data (Figs. 1E-F) and the KI67 immunofluorescence (Figs. 4A-B) is discussed and studied in detail in the next section.

**Fig. 4.**
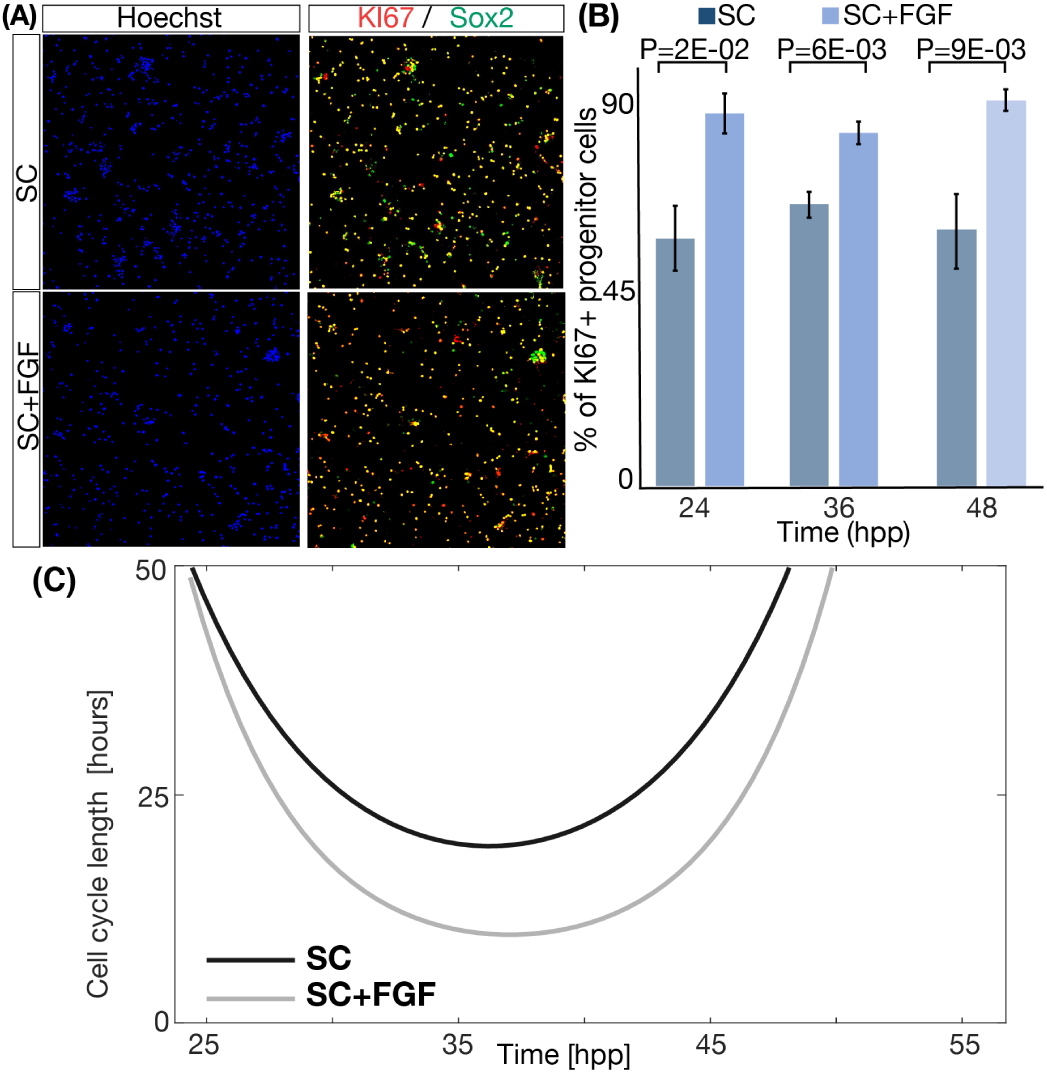
The growth fraction and the length of the cell cycle change in response to FGF2. (A) Example of cells stained with nuclei (blue), KI67 (red) and Sox2 (green) at 36 hpp. (B) Quantification of the percentage of progenitor cells that are actively cycling in both conditions and at three different time points. Columns represent the mean between independent repeats. Error bars represent the standard error or the mean. (C) Cell cycle prediction by the branching process formalism for the two different FGF2 concentrations tested.

The value of the cell cycle length obtained as output of Eq. 2 is plotted in Fig. 4C, showing an average value of *T* that is not constant: a continuous decrease in cell cycle length is followed by an increase at later time points, and the minimum values for SC (around *T* = 19 hours) and SC+FGF (around *T* = 10 hours) conditions occur around 36-37 hpp. These values are close to the values measured *in vivo* in Refs. (Gao et al., 2014; Beattie and Hippenmeyer, 2017), that report an average cell cycle length of 16-18 h in the temporal window corresponding to E11-E13.

### The branching process tool outperforms cumulative curve methods to monitor cell cycle dynamics

Interestingly, and despite showing the same trend of shortening *T* with FGF2, the absolute values of the cell cycle length predicted by the branching process formalism do not agree with the values obtained by the EdU cumulative experiments in Figs. 1E-F. This discrepancy in cell cycle dynamics and in the growth fraction (Fig. 4B) pointed us to study the potential source of conflict between the cumulative method and the Branching Process tool. To do that, we developed a numerical model of a generic differentiating stem cell population that simulates cycling progenitors that can either proliferate, differentiate, enter quiescence or apoptosis based on rates and probabilities provided by the user. Values of cell cycle length, mode of division, quiescence and death rate can be kept constant throughout the simulation, or can be set to change each time-step. Parameters are sampled from a gamma distribution to mimic intrinsic cell-to-cell variability. Details of the model are presented in the Supplementary Methods section. A scheme of the simulation framework is shown in Supplementary Figure S2.

A numerical analog of Edu is simulated computationally, in such a way that cells in S-phase are marked as labeled when EdU is present). Then, the number of progenitors, differentiated and EdU positive progenitors at each time point is used to calculate the average cell cycle length of the population using three widely used EdU based methods: single cumulative curve (**C1**) (Nowakowski et al., 1989), dual cumulative (**C2**) (Shibui et al., 1989), and the pulse-chase (**PC**) method (Weber et al., 2014). The cell cycle is also calculated using the branching process (**BP**) method (Míguez, 2015) (Eq. 2 in Supplementary Methods). A detailed description of each method and how it is applied in this context is illustrated in Supplementary Figure S3 and explained in the Supplementary Methods section. All predictions are then compared with the input value of *T* used for each simulation, to estimate the accuracy and reliability of each method.

The first scenario tested corresponds to homeostasis in the progenitor population (*pp* − *dd* = 0), constant value of *T* = 20 hours and no quiescent or apoptotic cells (*γ* = 1, ø_*P*_ = 0). These are the conditions defined by Nowakowski and coworkers when introducing originally the cumulative curve method (Nowakowski et al., 1989). Results of the analysis are plotted in Fig. 5A. Dots in Fig. 5B correspond to the prediction of the value of *T* for 10 independent simulations (crosses represent the average). We see that, for these particular settings, all four methods are able to predict the correct value of *T* (dashed line) within a 10% error margin, with both **PC** and **BP** performing slightly better than **C1** and **C2**. Importantly, when comparing the individual values for the 10 simulations predicted by single and double cumulative curve methods (**C1** and **C2**), there is a higher dispersion than in **PC** and **BP** methods. This means that a high number of repeats should be necessary to get an accurate value of *T*, and that the typical experimental design that involves only three independent repeats does not guarantee a correct estimation of the cell cycle. The same conclusions apply when considering growth of the population of progenitors, as in the case of RG reported here (*pp* − *dd* > 0, Supplementary Figure S4A).

**Fig. 5.**
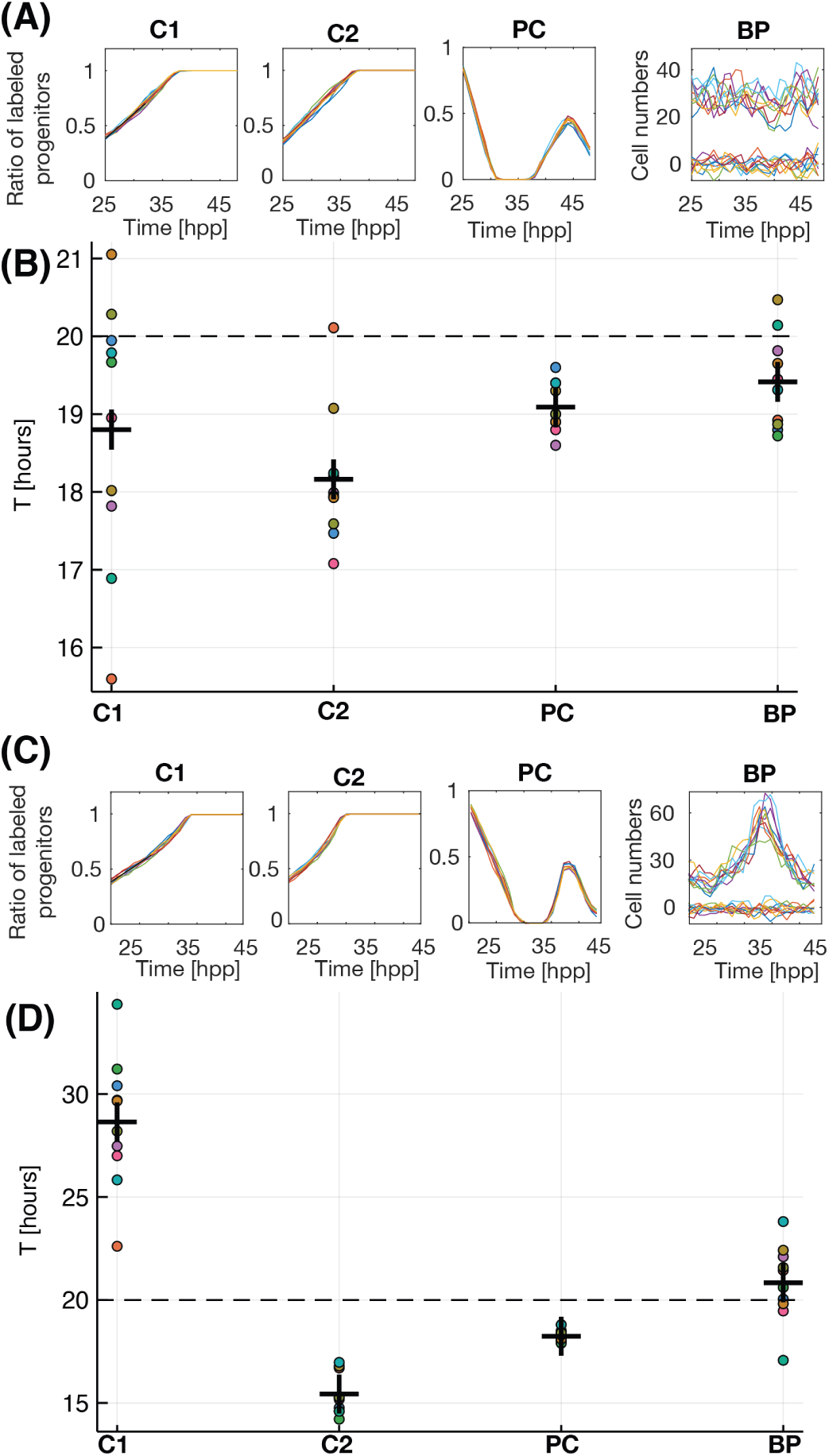
The branching process tool outperforms cumulative curve methods. Cumulative curves and quantification for Single Cumulative (**C1**), dual cumulative (**C2**), pulse-chase (**PC**) and Branching Process (**BP**) methods for 10 independent runs of the numerical model for conditions of (A-B) constant and (C-D) variable cell cycle length. Each color corresponds to the same simulation analyzed using each framework (see text). The cell cycle is also calculated using the branching process (**BP**) Dots correspond to single runs of the model, crosses show the average value for the 10 independent simulations.

Variable cell cycle dynamics has been reported in many developmental systems (Míguez, 2015; Saade et al., 2013; Le Dréau et al., 2014; Takahashi et al., 1995; Calegari and Huttner, 2003; Calegari et al., 2005; Dehay and Kennedy, 2007; Mairet-Coello et al., 2012; Roccio et al., 2013; Arai et al., 2011; Iulianella et al., 2008; Locker et al., 2006). Fig. 5C shows the output of the numerical model when a variable value of *T* is used as input (with an average value *T*= 20 hours). Fig. 5D plots the quantification of the cell cycle in these conditions. In this situation, **C1** predicts a much longer cell cycle that the average (49% error), while the **C2** predicts a shorter cell cycle (24% error). Interestingly both **PC** and **BP** return a value much closer to the correct average, with less than 10% error. Again, the variability of the single cumulative **SC** method (the one used in Figs. 1E-F and the most commonly used in the literature) is very high, making it unreliable when a small number of repeats are used (less than 10). Again, the same conclusions apply when considering conditions where the cell cycle changes while the population of progenitors is allowed to grow (*pp* − *dd* > 0, Supplementary Figure S4B).

The balance between differentiative and proliferative divisions has been shown to also change overtime in many developmental systems (Saade et al., 2013; Míguez, 2015). For instance, during motorneuron generation, the rate of differentiation changes rapidly due to a sudden switch in Shh levels (Saade et al., 2013). We show here that even *in vitro*, with cells growing in constant controlled conditions, the mode of divisions is highly non-constant (Fig. 3A). When we set a variable differentiation rate in our simulations, we observe that again both single **SC** and dual **DC** cumulative methods fail and show high dispersion between independent samples (Supplementary Figure S4C). The same occurs when both mode and rate of division are allowed to change simultaneously (Supplementary Figure S4D). In these more realistic conditions closer to our experimental findings (variable mode and rate of division), the branching process equation predicts a value that is closer to the one used in the simulations, and the variability between samples is highly reduced.

In conclusion, these results show that methods based on cumulative curve labeling are not suitable when proliferation and/or differentiation rates are not constant. This, together with the reported effect of BrdU and analogs in lengthening the cell cycle (Levkoff et al., 2008), and the high dispersion when comparing sets of cells growing at the same exact conditions, could explain the discrepancy values of the cell cycle reported in Figs. 1E-F and Fig. 4C. In addition, the error in the growth fraction measured in Figs. 1E, F versus Fig. 4A can be due to the same problems. Both Pulse-Chase **PC** and Branching Process **BP** perform well, while the branching formalism has the advantage of providing temporal resolution, as well as accurate values of the average mode of division during the experiment.

### Values from the branching process analysis are able to reproduce the experimental data

To test if the values provided by the branching process formalism are correct, we take advantage of the same numerical model of the differentiating stem cell population introduced previously. Now, the model is informed with the values of initial number of cells as in the experiments, the prediction of *T* and *pp* − *dd* predicted by the branching process (Fig. 3A and Fig. 4C), and the growth fraction *γ* and apoptosis measured in the previous sections (Figs. 4B, and Supp Fig. 1B).

Results are plotted in 6A-B, where we plot the prediction for number of progenitors and differentiated cells (thin green and red lines, respectively) for 30 independent simulations. Comparison with the fitting of the experimental data for progenitors and differentiated cells (thick green and red lines, respectively) show a good agreement in both conditions, suggesting that the branching equations are able to predict the correct average mode and rate of division of RG *in vitro*.

## Discussion

A detailed analysis of the dynamics of vertebrate neurogenesis involves a careful characterization of the features that regulate the dynamics of proliferation and differentiation of RG during the generation of the mammalian cortex. One of its most striking features is the fact that RG are restricted to an asymmetric mode of division *in vivo*, as oppose to a more probabilistic scenario observed in other developmental systems (Saade et al., 2013; Míguez, 2015; He et al., 2012; Chen et al., 2012; Clayton et al., 2007; Teixeira et al., 2013; Klein et al., 2010; Snippert et al., 2010). FGF2 has been shown to facilitate the expansion of RG *in vitro* cultures, but the details of this process have not been studied. Our quantitative characterization of the effects of FGF2 show a multiple effect in the growth fraction (Fig. 4B), the mode of division (Fig. 3A) and in the length of the cell cycle (Fig. 4C).

The overall influence of each of these effects in the expansion potential of the RG culture can be assessed using our numerical model. To do that, we inform the simulations with the experimental values for SC, and quantify the increase in the number of cycling progenitors after 22 hours (as a measure of the potential of the culture to expand in size). Next, we substitute each of the predictions for cell cycle length, growth fraction and differentiation rate predicted for the SC+FGF2 conditions, individually or in combination. The increase in cycling progenitors for 30 independent numerical simulations for each condition is shown in Fig. 6E. Surprisingly, the analysis suggests that the most influential feature is not the differentiation rate or the growth fraction, but the change in cell cycle length. The change in growth fraction or differentiation rate do not significantly impact the culture in terms of cycling progenitors (1 % and 9%, respectively), but when combined with the effect on the cell cycle, they can increase the expansion by an additional 25%.

**Fig. 6.**
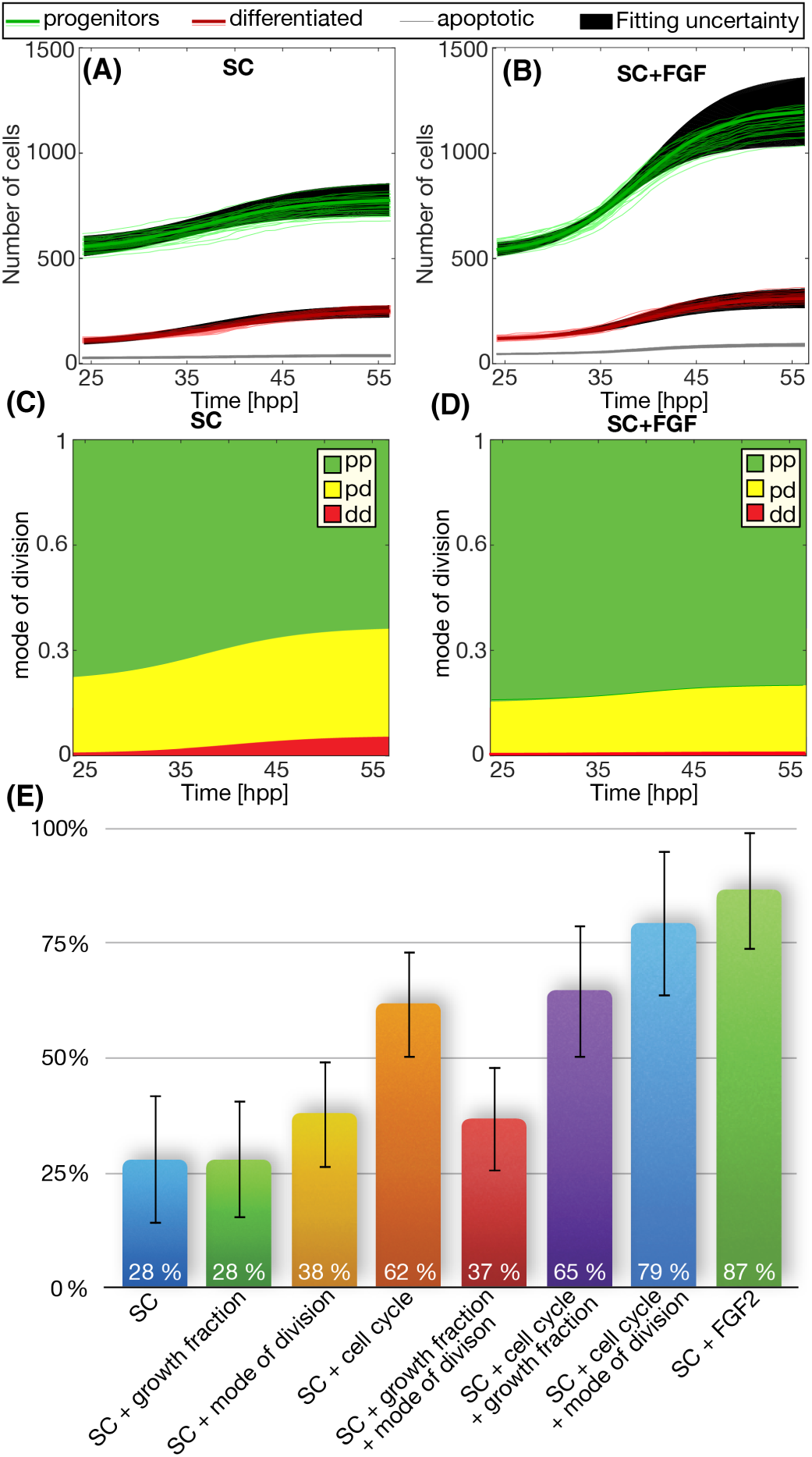
Values derived using the branching process formalism reproduce the correct dynamics observed experimentally. (A-B) Numerical simulations (light red and green lines) for both conditions using the parameters of mode and rate of division predicted by the equations of the branching process. Thick lines correspond to the sigmoidal fitting of the experimental data in Fig. 2. Black regions mark the uncertainty that result from the fitting. (E) Changes in the population of cycling progenitors due to each of the three effects of FGF2 alone or in combination.

Several authors propose that the mode of division depends on the distribution of cell fate determinants during mitosis, the orientation of the spindle or the inheritance of the primary cilium or the different centrosomes (Taverna et al., 2014). It is possible that the apical-basal polarized structure of the RG, or their organization and orientation of the radial processes along the stratified neuroephitelium results in asymmetric inheritance of these cell fate regulators (Taverna et al., 2014). The loss of these polarizing features provided by the niche when cells are cultured *in vitro* may result in a probabilistic scenario where the fate of the two daughter cells is independent of each other and all of the 3 modes of divisions are possible, similarly to neuronal progenitor cells and other developmental systems (Saade et al., 2013; Míguez, 2015; He et al., 2012; Chen et al., 2012; Clayton et al., 2007; Teixeira et al., 2013; Klein et al., 2010; Snippert et al., 2010). In fact, early studies in the mouse neocortex suggest that the model that fits best the clone distribution assumes that the fate of the daughter cells is independent of each other (Cai et al., 2002). In this situation, the branching process framework is able to estimate the rates of each of the three modes of division (Míguez, 2015). This prediction for the case of RG in culture is shown in Fig. 6C-D, where we can see that the predominant mode of division is *pp* (green). This symmetric mode of division is even more probable in conditions of SC+FGF, to the expenses of a reduction in *pd* and *dd*.

A detailed analysis of the dynamics of vertebrate neurogenesis involves a careful characterization of the rate of division. The most direct method to measure the cell cycle length requires to monitor the time between consecutive mitotic evens at single cell resolution (Sigal et al., 2006). Unfortunately, due to the high degree of variability, many cells in a population need to be sampled, segmented and tracked simultaneously to obtain an accurate value, even when dealing with clonal samples (Sandler et al., 2015). Therefore, the most used approach is the use of thymidine analogs, but this approach has several drawbacks: it can be toxic and mutagenic (Duque and Gorfinkiel, 2016) and affect the normal dynamics of cell proliferation (Levkoff et al., 2008) by lengthening the cell cycle. In addition, choosing the correct mathematical analysis and interpretation of the experimental data is not straight-forward (Johansson et al., 1999). Authors have proposed several approaches, such as linear (Begg et al., 1985; Hoyer et al., 1994), nonlinear fitting (Johansson et al., 1994; Weber et al., 2014), or the use of deterministic (Lee and Perelson, 2008) or stochastic models (Zilman et al., 2010). Depending on the method used, the same input data results in quite different predictions for the average duration of the cell cycle (Ritter et al., 1992). Due to these limitations, BrdU and analogs have been referred as “one of the most misused techniques in neuroscience” (Taupin, 2007).

Our results shown that methods based on cumulative incorporation of thymidine analogs perform well in conditions of constant proliferation and differentiation, but they are not designed to study systems where the cell cycle changes overtime, which is is potentially the case in many developmental systems. In these conditions, the Branching Process formalism and the Pulse-Chase outperform cumulative curve methods. On the other hand, the Pulse-Chase method requires experiments that are longer than the cell cycle length, so an estimation of the value of the cell cycle has to be known beforehand. In addition, the toxic effect of the labeling agent for such long periods of time may affect strongly the normal cell cycle progression, probably by enlarging its real value (Levkoff et al., 2008; Duque and Gorfinkiel, 2016). A clear advantage of the Branching Process is that it does not involve manipulation of the samples before fixation, so there is no interference with the normal progression of the cell cycle. In addition, the Branching Process formalism also provides the correct value of *T* with temporal resolution, and the measurement of the average differentiation rate, (also with temporal resolution).

Several studies have shown that the length of G1 phase increases progressively when neurogenesis starts, resulting in a overall increase of the cell cycle (Takahashi et al., 1995; Calegari and Huttner, 2003; Calegari et al., 2005; Dehay and Kennedy, 2007; Mairet-Coello et al., 2012; Roccio et al., 2013). Alternatively, others studies show that the cell cycle length is shorter in neurogenic divisions, compared to proliferative divisions (Arai et al., 2011; Saade et al., 2013; Le Dréau et al., 2014; Iulianella et al., 2008; Locker et al., 2006), due mainly to a shortening in S-phase. Our results show that FGF promotes pp divisions and shortens cell cycle, consistent with the hypothesis that proliferative divisions have a shorter cell cycle, maybe via a shortening of G1-phase (similarly to insulin-like growth factor (Mairet-Coello et al., 2009; Hodge et al., 2004)). A careful characterization of how FGF2 affects each phase of the cell cycle it is far from the scope of this contribution.

## CONCLUSION

The culture and differentiation of RG cells *in vitro* provides a very good framework to study basic features that orchestrate the formation of the mammalian neocortex. In brief, the system provides a well controlled environment where the effect of signaling molecules and other conditions can be tested reliably, while providing easier manipulation and imaging compared to studies performed *in vivo*. We use this framework to study the features that promote the expansion of RG in culture driven by FGF2. Our combined experimental/computational/theoretical approach can be also used to test the effect of other signaling networks by quantifying the cell cycle and mode of division after ligand stimulation or small molecule inhibition, after a comparison with a control culture.

## Materials and Methods

### Preparation and culture of dissociated mouse cortical RG

Cells were obtained from mouse embryos of the C57 BL/6JRCC line at E11/E11.5, following standard methods described previously (see Ref. (Hilgenberg and Smith, 2007)). The initial time point is labeled as 0 hours post plating (hpp) and it is used as the reference point for our experiments. Briefly, after careful removing of the meninges, the cortex is isolated and placed in Hank’s Buffered Salt Solution free of Ca2+ and Mg2+ (HBSS, ThermoFisher 14185). Next, samples are mechanically disgregated using Pasteur pipettes and plated in coverslips treated with Nitric Acid and Fibronectin at 10 µg/ml (Fisher Scientific; 15602707) to facilitate cell adhesion. Cells are plated at constant density (250000 cells in each M24 well) for all experiments in Neurobasal medium without L-glutamine (ThermoFisher 21103-049), Glutamax (ThermoFisher 35050-038), B-27 (ThermoFisher 17504-044) Penicillin, Streptomycin and Antimicotic (concentrations standard for cell culture). Media is complemented with 0.02 ng/*µ*l of recombinant murine EGF (PeproTech 315-09, lot number 0517179-1) and 0.02 ng/*µ*l of human FGF basic (PreproTech 100-18B, lot number 0311706-1). This culture media is referred as *standard culture* (SC) conditions in our study. Cells are allowed to rest a full day in the incubator to recover the dissection process. 24 hpp, the culture media is changed with fresh SC media, or to SC media complemented with additional human FGF basic to a final concentration of 0.06 ng/*µ*). This culture conditions are labeled as SC+FGF in this study. All experimental protocols were in accordance with the guidelines of the European Communities Directive (2012/63UE) and the actual Spanish legislation (RD 53/2013).

### Immunofluorescence

Cells are fixed for 20 minutes at Room Temperature (RT) with 4% paraformaldehyde and washed twice for 5 minutes with Phosphate Buffer Saline 1X (PBS). Fixed cells are incubated with the permeabilization solution composed of Triton x-100 (ChemSupply 9002-93-1) at 0.6% in PBS 1X for 20 minutes at RT. Next, cells are washed 3 times with PBS and blocking solution is added (Bovine Serum Albumin, BSA. Sigma; A7906) at 3% in PBS for at least 30 minutes. Later, cells are incubated with primary antibodies dissolved in the blocking solution overnight at 4°C. The next day, cells are washed with PBS 3-4 times for 5 minutes, and they are incubated with secondary antibodies in the blocking solution for 45 minutes at RT, protected from light. Next, secondary antibodies are washed out (PBS 3-4 times for 5 minutes), and nuclei is stained with Hoechst 3342 (1/2000, ThermoFisher 1399) dissolved in PBS for 5 minutes at RT. Finally, cells are washed in PBS, double distilled water, and ethanol at 70%. Cover-slips are finally mounted with Fluoromount G (Southern Biotechnology Associates, Inc, Birmingham, Alabama 0100-01) on microscope glass slides. Primary antibodies used are: anti-Sox2 (1/2000, Gene-Tex GTX124477), anti-Map2 (1/200, Santa Cruz Biotechnology sc-74421); anti-Pax6 (1/1000, BioLegend B244573); anti-Cleaved Caspase 3 (1/1000, Cell Signaling 9661); and anti-KI67 (1/200, ThermoFisher 14-5698-82). Secondary antibodies used are: anti-Rabbit 488 (1/1000, ThermoFisher A-21206), anti-Mouse 555 (1/1000, ThermoFisher A-21137) and anti-Rat 555 (1/1000, ThermoFisher A-21434).

### Statistical and Data analysis

One way ANOVA test is used to measure statistical significance between different time points. Cell cycle values in Fig. 1E-F are obtained after linear regression of the four first data points. Rates of quiescence in Fig. 4B are obtained from the mean value of the four last points. Slope error is calculated doing a linear fitting with values of the average plus standard error and another one with values of the average minus standard error to get the difference in the slope between these two values. Quiescence error is the standard error of the four last points, and the *T* error is derived from the error propagation of the previous values. Three-parameter sigmoidal fitting is used to fit data from Fig. 2B-C. Black regions in Figs. 6A-B mark the uncertainty derived from the fitting, calculated from the difference between the result of the fitting using as values the mean plus the standard error, and the result of the fitting using as values the mean minus the standard error (with the same values of the parameters). Sample size for all experiments is at least 4. Unless specified, errorbars represent the standard error of the mean, calculated using error propagation. All curve fitting and statistical analysis are performed using Matlab^©^ (The Mathworks^©^, Natick, MA) and Julia programming language (Statistics package).

### EdU cumulative curve

Cumulative curve of the thymidine analog 5-Ethynyl-2’-deoxyUridine (EdU) incorporation is performed using Click-iT™ Plus EdU Alexa Fluor™ 647 Imaging Kit (ThermoFisher; C10640). Briefly, EdU was added around 24 hpp at 2 *µ*M. Cells are then fixed at increasing times of EdU exposition. Staining of EdU positive cell is performed based on previously published protocols (Harrison et al., 2018). Next, immunostaining against Sox2 is used as standard marker for RG progenitors (Beattie and Hippenmeyer, 2017). Later, the number of cells positive for both Sox2 and EdU is quantified using automated image processing. To calculate the cell cycle length, the percentage of progenitor cells that have incorporated EdU is plotted against the hours of EdU incorporation. The saturation value at long incubation times is used to calculate the growth fraction *γ*. This value is then used to calculate the average cell cycle using linear regression at short EdU accumulation times (see figure 1).

### EdU pulse-and-chase experiments

Cells are exposed to a short pulse of 30 minutes of EdU at 36 hpp. “Pulse” points are fixed at this time point. “Chase” points are washed three times with fresh medium and are fixed 15 hours after the “Pulse” time point. The number of EdU positive/Sox2 positive cells is quantified in both “Pulse” and “Chase” time points for both conditions using automated image processing.

## Acknowledgements

We thank professor Francisco Wandosell for multiple discussions and invaluable input at all stages of this work.

## Contribution

MLT performed research and wrote the manuscript, NPC, performed research, DGM designed research, performed research and wrote the manuscript.

## Data availability

Code for numerical simulations model is available as supplementary material

## Supplementary

### Image acquisition and analysis

Samples are imaged in a confocal microscope AR1+ of high speed in acquisition and sensibility coupled to an inverted microscope model Eclipse Ti-E (Nikon) with a 20X objective and a resolution of 1024×1024 pixels. The field of view is set to 0.6 mm × 0.6 mm. In brief, image processing and analysis (performed in Fiji (Schindelin et al., 2012)) is based on the segmentation of nuclei and the classification of each cell as progenitor, differentiated neuron, quiescence or apoptotic based on the intensity of the fluorescence staining of each marker. A large number of cells (around 10^5^ cells) is processed for each data point to minimize the effect of variability and heterogeneity of the samples. The sequence of processing algorithms and filters is as follows.

1. Definition of the Kernel Radius (*KR*) that sets the size of the region used for calculations and filter processing. Several *KR* sizes were tested (values from 1 to 5 pixels). The final *KR* was fixed as 2.5.
2. A local thresholding is applied to remove background based on the median intensity as cutoff value (radius= 8x*KR*).
3. To remove breaks and holes inside the objects generated by the previous filter, the following sequence of filters is applied to enhance the definition of the boundaries of each object: Gaussian Blur filter, Maximum Filter, Median filter and Unsharp Mask filter (radius = KR).
4. The resulting image is binarized using the median value as threshold.
5. Euclidean Distance Map (EDT) is performed in the binary image to generate seeds that are used by a flood fill algorithm to define the boundaries of each object (Kang et al., 2010).
6. Finally, all objects are fitted to ellipses for posterior analysis. Ellipses smaller than 4 × *π* × *KR*^2^ are discarded from the analysis.

The specific features of each staining requires a different set of processing filters to enhance signal for each channel.

1. Map2: Double sequential thresholding to extract foreground information (cut-off 1 = mean, cutoff 2: median); morphological opening to remove neurons fibers (structuring element: lines at different angle with a length of 2x*KR*); Gaussian filter to remove noise (radius=KR).
2. Sox2: Double sequential thresholding to extract foreground information (cuttof 1, = mean, cutoff 2: median); morphological opening to select only nuclei with minimal size (structuring element: circumference of radius equal to 2x*KR*); Gaussian filter to remove noise (radius=KR).
3. EdU: Single thresholding to extract foreground information (cutoff = median); morphological opening (structuring element: circumference of radius equal to 2x*KR*); Gaussian filter to remove noise (radius=KR).
4. Pax6: Single thresholding to extract foreground information (cutoff = mean); morphological opening (structuring element: circumference of radius equal to 2x*KR*); Gaussian filter to remove noise (radius=KR).
5. Cleaved Caspase-3: Double sequential thresholding to extract foreground information (cutoff 1= mean, cutoff 2 = mean + plus standard deviation); morphological opening to select only nuclei with minimal size (structuring element: circumference of radius equal to 2x*KR*); Gaussian filter to remove noise (radius=KR).
6. KI67: Single thresholding to extract foreground information (cutoff = mean); morphological opening (structuring element: circumference of radius equal to 2x*KR*); Gaussian filter to remove noise (radius=KR).

Finally, the identity of each ellipse is established based on the number of pixels above threshold in each channel. For the MAP-2, this area was set to at least 15%, and 1% for the rest. A subset of cells are both Sox2- and Map2-, and have a nucleus that is much larger that Sox2+ or Map2+. Since these cells are not RG or differentiated neurons, they are not taken into account in the study.

### Numerical simulations of cell populations

We developed an *in silico* phenomenological model of the culture of cells as numerical entities that proliferate, differentiate enter quiescence or apoptosis according to probabilities established by the user. Each cell has the following features: length of its cell cycle (T), current phase of cell cycle, time since birth (age), and fate (progenitor, quiescent, differentiated or apoptotic). These features are updated for each cell at each time point, since they can change due to events such as cell division, changes in the culture media. Values of average mode of division, average cell cycle length of the population and percentage of cycling progenitors, percentage of cell undergoing apoptosis, are set by the user. To mimic the inherent cell-to-cell variability and intrinsic noise in a clonal population (León et al., 2004), the value for each parameter is obtained from a gamma distribution with mean defined by the user and standard deviation of 30% of the mean (other values from 0% to 50% provide similar results).

A scheme of how the population is defined and develops overtime is shown in Supp Fig 2. Parameters of the simulation are: the number of initial cells *m*, the average cell cycle *T* at each time point (defined as *T* = ∑*T* ^*i*^/*n*, being *n* the number of cells at time *t*), the fraction of cycling progenitors (or growth fraction) *γ*, the rate of apoptosis of progenitors ø_*P*_, and the length of the experiment *t*_*end*_. The age of each cell is defined as the time since its birth, and the type corresponds to its characteristic as progenitors (*P*, cycling cells), differentiated (*D*, non cycling cells), quiescent (*Q*, non cycling progenitors) and apoptotic (dying cells).

The simulation takes palace as follows: an initial set of un-synchronized progenitors cells are allowed to cycle following the different phases of the cell cycle: from *G*_1_ to *S* to *G*_2_ to finally *M* phase. Upon division, the two resulting daughter cells either remain as progenitors (*pp* division), they become terminally differentiated cells and stop cycling (*dd* division), or one remains as progenitor while the other differentiates (*pd* division). For simplicity, the cell cycle is divided into just three main steps of equal length: *G*_1_, followed by *S* and finally followed by *G*_2_ + *M*. (*T* = *T*_*G*1_ + *T*_*S*_ + *T*_*G*2*M*_). Changes in the cell cycle length affect all phases of the cell cycle identically (simulations where the phases are of different length and that changes affecting differently different phases of the cell cycle show equivalent results).

### Branching Process formalism

Our lab has developed a method to measure the dynamics of proliferation and differentiation that do not depends on thymidine cumulative labeling. Instead, it uses a branching process formalism to obtain analytical equations that provide the average values of proliferation and differentiation of the population based only on the numbers of proliferative, differentiated, quiescent and apoptotic cells at different times points. A scheme of the method is shown in Supp Fig 3D, and an example of its experimental implementation can be found in Ref. (Míguez, 2015).

To obtain these values, samples are allowed to develop without interfering with the normal dynamics of the cells, and then are fixed at different developmental times. After fixation, the amount of cells in each state is quantified by antibody staining to distinguish progenitors (*P*) (Graham et al., 2003), differentiated (*D*) (Míguez, 2013), and the number of progenitors undergoing apoptosis (ø_*P*_) (Blanchard et al., 2010). The growth fraction *γ* is obtained using double immuno-labeling against Sox2 and KI67 (Scholzen and Gerdes, 2000).

These values (quantified using the automated quantification described in the Methods Section) are used as input of the following equations for the mode and rate of division, which correspond to a generalization of Equations presented in Ref. (Míguez, 2015) updated to account for a potential reduction of the progenitor pool:

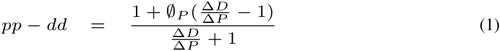

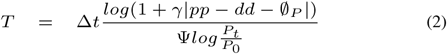

where *pp* and *dd* correspond to the rate of symmetric proliferative and differentiative divisions, respectively. Δ*P* = *P*_*t*_ − *P*_0_ and Δ*D* = *D*_*t*_ − *D*_0_ correspond to the number of progenitors and differentiated cells generated in a given window of time Δ*t* = *t* − *t*_0_. The value *pp* − *dd* goes from 1 (all divisions are symmetric proliferative) to −1 (all divisions are symmetric differentiative). The value, *pp* − *dd* = 0 corresponds to maintenance of the progenitor pool, either via asymmetric *pd* divisions of via balance between symmetric proliferative and differentiative divisions (the model cannot distinguish between these two scenarios, since they are mathematically equivalent). Ψ takes the value of 1 when 1 ⩽ *pp* − *dd* < 0, while for values between 0 ⩽ *pp* − *dd* ⩽ −1 takes the form Ψ = (0.9|*pp* − *dd*| − 1). ø_*P*_ is the rate of cell death of the progenitors pool, obtained using double immuno-labeling against Sox2 and Cleaved Caspase3 (see Supp Fig 1B). This reduced value of apoptosis rate (assuming that most cell death occurs via apoptosis) is consistent with estimations from *in vivo* experiments (Cai et al., 2002).

### Simulations of Cell Cycle determination methods

The previous model is then adapted to perform a computational analog of one or two thymidine compounds. At any time in the simulation, EdU can be added to the cells, so cells undergoing S-phase will be labeled as “positive”, and will remain as positive throughout the rest of the simulation. The input parameters of the model are varied to simulate different dynamics of a population of cells in different conditions, in terms of quiescence, apoptosis, cell cycle length and differentiation rate. For each condition tested, we will perform four measurements of the cell cycle based on the following methodologies:

#### Cumulative Curve method

This technique has been extensively used both in *in vitro* and *in vivo* situations to quantify the rate of cells in the population entering S-phase (Martinez-Morales et al., 2010; Le Dréau et al., 2014). A scheme of the method is shown in Supp Fig 3A. In brief, a nucleoside analog is added to several identical samples that are fixed and stained at different times. Labeled cells in all samples are quantified using microscopy or flow-cytometry. The ratio of progenitor cells that are labeled for each sample is plotted, and the values corresponding to the cell cycle length *T* are obtained from the slope of a linear regression fitting of the data at short exposure times. In addition, the fraction of cycling progenitor cells *γ*, or growth fraction, can be estimated from the rate of labeled cells after long exposure times. This method, when combined with dyes to measure DNA content can be used to determined the length of the different phases of the cell cycle (Dolbeare and Selden, 1994).

#### Dual Cumulative Curve Method

This method combines dual staining with thymidine analogs (Salic and Mitchison, 2008). It also provides the possibility of fixing all samples simultaneously to ensure that quantification is performed always at the same developmental time. In addition, it can also provide some positional information of regions in a given tissue where cells cycle at different rates (Shibui et al., 1989; Bradford and Clarke, 2011). On the other hand, it requires a more complex experimental design, and it may also result in increased toxicity. In addition, it does not provide information about the growth fraction. The method (Supp Fig 3B) involves a first labelling agent administered to all samples simultaneously, and a second agent administered at different time points. All samples are collected at the same time, and they are stained for both labelling agents. The amount or cells that are double positive overtime for the two different thymine analogs is plotted, and the average length of T and *T*_*S*_ can be obtained using linear or nonlinear regression (some corrections regarding the potential differential incorporation of both agents are required).

#### Pulse-Chase method

Both previous methods rely on long term exposure of the samples to nucleoside analogs, which can result in toxicity effects. Alternative, a short pulse can be also applied (Weber et al., 2014) to label only cells that where in S-phase at a given time. Then, the population of positive cells is “chased” in the different samples by fixing and staining at different times. Several variations of this method have been developed. A commonly used technique is to stain cells in mitosis (using immunofluorescence against phospho-histone-3), or using a second thymine analog in S-phase to chase cells that have re-entered in a new S-phase. A scheme of the method is shown in Supp Fig 3C.

The ratio of double positive cells in the different samples is plotted overtime, and the average value of *T* corresponds to the time between the pulse and the maximum of double positive cells in the population. The slope of the curve at shorter time scales can be used to calculate the length of S-phase. Measurements of the cell cycle using this methods requires significantly longer experiments than the two previous methods.

#### Branching process Method

The number of cells and their fate as progenitors, differentiated, quiescent or apoptotic is recorded at each time point during the simulation. These values are then used as input of the branching process equation 2 described briefly in other subsection of the Methods section. The average value is then plotted for each condition tested.

**Fig. 7.**
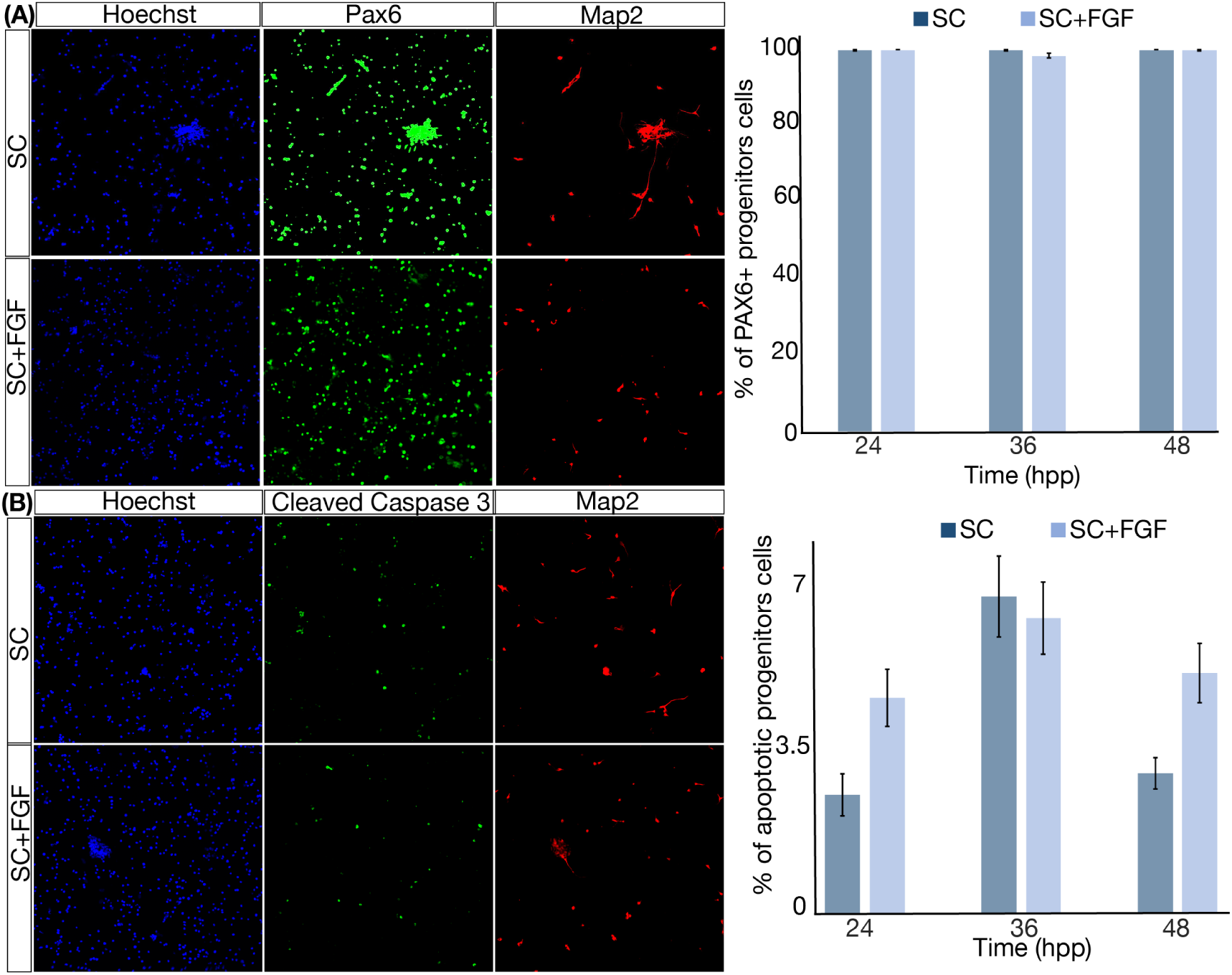
Supplementary Figure 1: Experiments to obtain the apoptosis rate and the amount of NEP. (A) Example of cells stained with nuclei marker (blue), Pax6 (green), and Map2 (red) at 36 hpp. Quantification of the percentage of progenitors that are Pax6 positive for all conditions and three time points. Columns represent the mean between independent repeats. (B) Example of cells stained with nuclei (blue), cleaved Caspase3 (green), and Map2 (red) at 36 hpp. Quantification of the percentage of progenitor cells that show positive staining for Caspase3 for both conditions and at three different time points. This low value of apoptosis rate is consistent with estimations from *in vivo* experiments (Cai et al., 2002). Error bars represent the standard error or the mean.

**Fig. 8.**
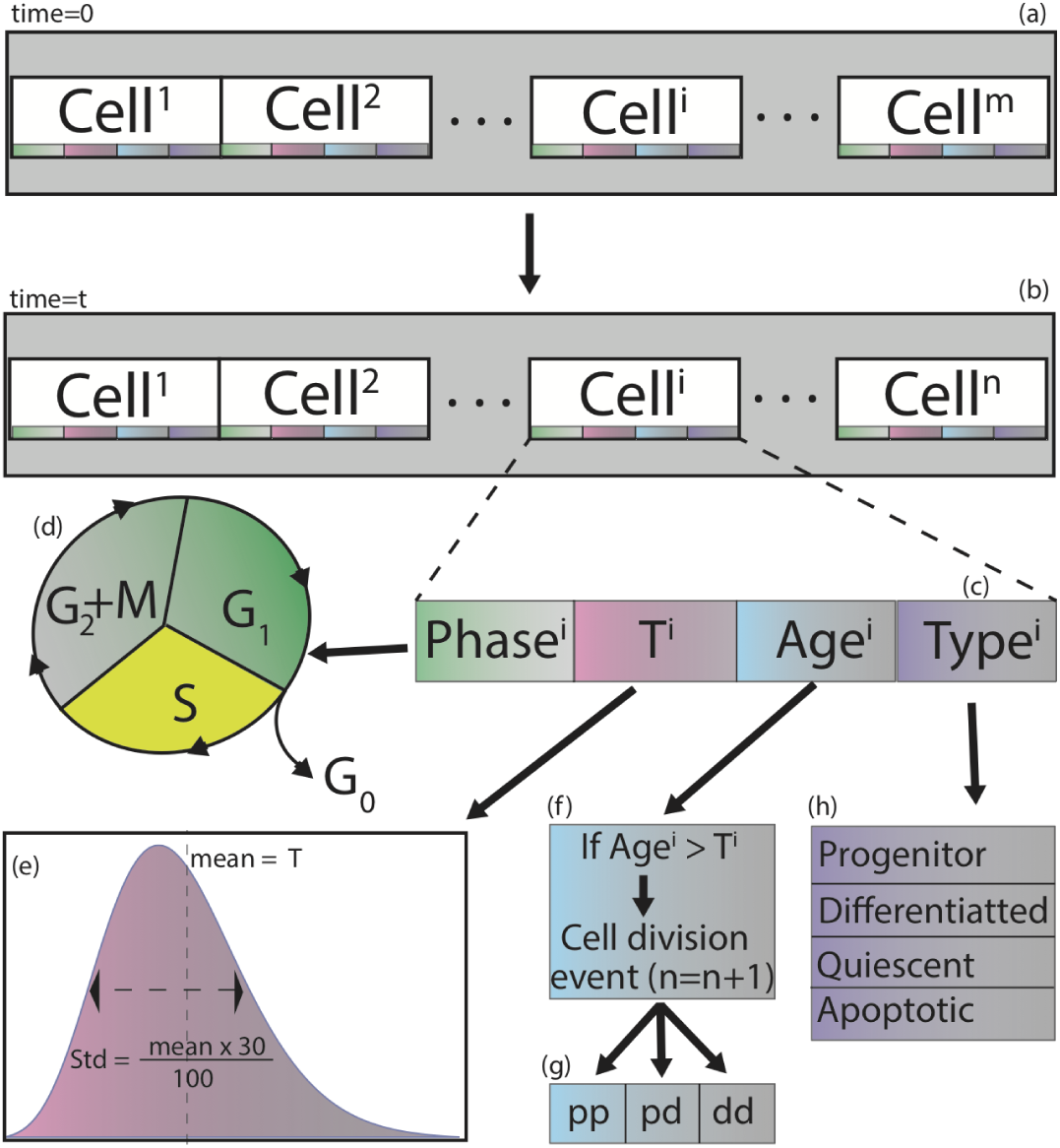
Supplementary Figure 2: Scheme of the simulation of the population model. In brief: (a) an initial number *m* of un-syncronized progenitor cells proliferates and differentiates. (b) At any given time *t*, each cell *i* in the population of *n* cells is characterised by four parameters (c): phase, T, age and Type. (d) Cells cycle in their phase from *G*_1_ to *S* to *G*_2_ + *M*. When a given cell *i* reaches the end of *G*_2_ + *M*, a division event takes place, with three different outcomes (g): *pp, pd* or *dd* division. In the presence of a labelling agent, cells incorporate it only during S-phase, and become labeled as “positive” (showed in yellow). Depending on their type, cells are sorted into 4 groups (h): progenitors, differentiated, quiescent and apoptotic.

**Fig. 9.**
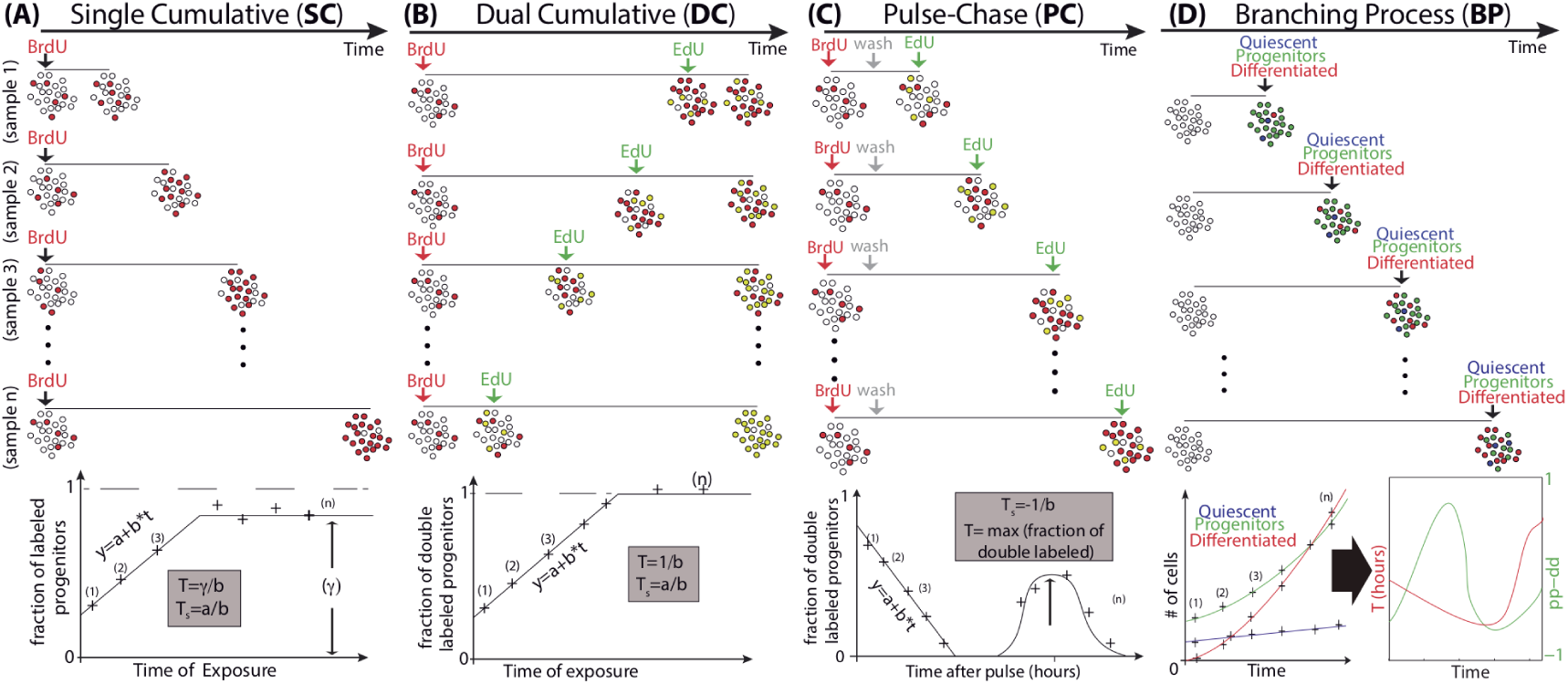
Supplementary Figure 3: Scheme of methods tested to measure cell cycle length. (A) Cumulative curve: A thymidine analog is added to all samples simultaneously. Samples are fixed at different times and stained for quantification. Linear fitting of the rate of labelling is used to determine the average *T* and *γ*_*P*_. (B) Dual Cumulative: The first thymidine analog (red) is administered to all samples simultaneously. The second thymine analog (green) is administered at different times. All samples are then fixed simultaneously. Quantification of all double positive cells (yellow) is plotted against exposure time. This method does not provide an estimation of the growth fraction. (C) Pulse-chase: A short pulse of a first nucleoside analog is added to all samples simultaneously. A second nucleoside analog is added at different times, and the samples are fixed and stained immediately after. The amount of double positive cells is plotted overtime. (D) Branching process: Cells are fixed at different times and stained with antibodies to distinguish progenitors, differentiated, quiescent and apoptotic cells. The resulting numbers are used to inform the equations 1-2, that will give us the values of the average rate and mode of division overtime.

**Fig. 10.**
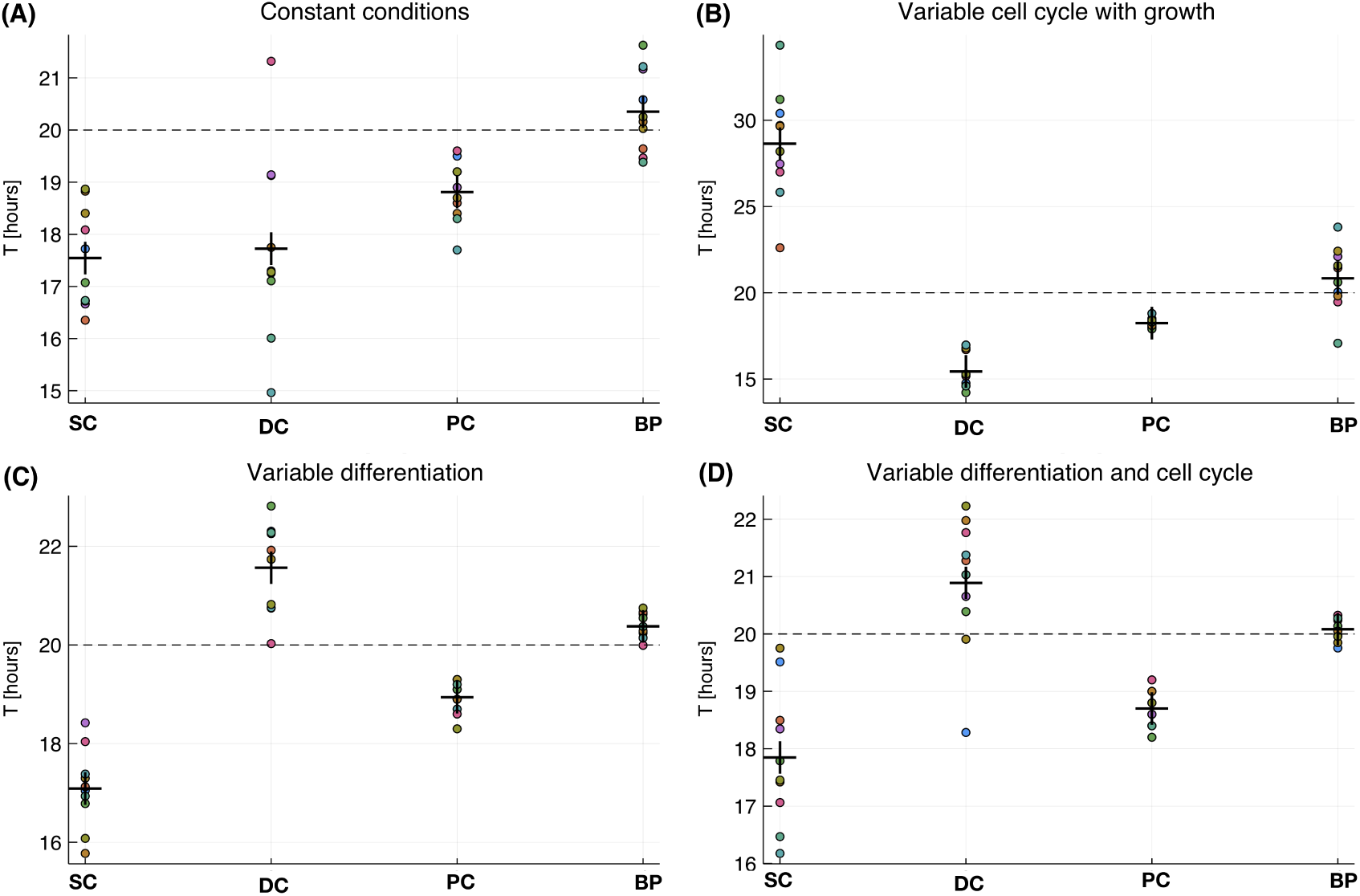
Supplementary Figure 4: Values of cell cycle predicted for different conditions of growth and differentiation of the culture. Dots correspond to independent simulations. Crosses represent the average between 10 simulations. Dashed horizontal line corresponds to the average value of *T* used in the simulations (20 hours). Shorter distance between crosses and dashed line represent better performance of the method. Lower dispersion between dots in each method represents better accuracy. (A) Predicted value of *T* by each method in conditions of constant mode and rate of division, but for values of increased in the population of progenitors (*pp* − *dd* > 0). (B) Predicted value of *T* by each method in conditions where the cell cycle is set to decrease and then increase. (C) Predicted value of *T* by each method in conditions where the differentiation is increasing monotonically during the simulation. (D) Predicted value of *T* by each method in conditions where both cell cycle and differentiation rate are set to change during the simulation.

